# A circRNA-derived IRES-like element enables programmable translational activation of the endogenous chaperone HSP90

**DOI:** 10.64898/2026.10.05.756760

**Authors:** Guy Aoun, Andrea Vandelli, Riccardo Delli Ponti, Stefano Gustincich, Elias Bechara, Laura Broglia, Gian Gaetano Tartaglia

## Abstract

Circular RNAs (circRNAs) are covalently closed transcripts that lack a 5′ cap, consequently their translation depends on cap-independent initiation, in most cases through Internal Ribosome Entry Sites (IRESs). Whereas IRES-mediated translation has been studied almost exclusively *in cis*, the use of IRES-containing circRNAs to activate translation of an independent mRNA *in trans* has only recently begun to be explored. Here, we characterize *SynFarsa*, a synthetic circRNA-derived IRES-like element that promotes cap-independent translation in mouse and human cells. We engineered *SynFarsa* into a *trans*-acting regulator by fusing it to a short antisense sequence complementary to a selected region of a target mRNA. This reproduces the architecture of a circular SINEUP, in which IRES activity promotes translation *in trans* of a transcript recognized through antisense complementarity. Directing this construct to the *hsp90* 3′UTR increased HSP90 protein levels approximately 1.6-fold without altering *hsp90* mRNA abundance, whereas the *hsp70* 3′UTR and the 5′UTRs of both transcripts were not permissive, indicating that activity depends strongly on target site selection. In a FUS^P525L^ neuroblastoma model of Amyotrophic Lateral Sclerosis, this increase in HSP90 was accompanied by a reduction in the number and size of cytoplasmic FUS aggregates. *SynFarsa* thus provides a circRNA-derived effector domain for targeted translational upregulation of an endogenous protein.

## Introduction

Circular RNAs (circRNA) are single-stranded RNA molecules characterized by a covalently closed structure lacking the free 5′ and 3′ ends. They are generated through back-splicing, a non-canonical splicing event in which a downstream splice site donor is joined to an upstream splice site acceptor, producing a circular transcript (1,2). Their circular structure provides increased resistance to exonucleolytic degradation and contributes to their stability compared with many linear RNA molecules (3,4). circRNA biogenesis is influenced by the organization of the surrounding genomic region, including complementary or inverted intronic sequences and by RNA-binding proteins (RBPs) that bind the intronic regions flanking circularizing exons and thereby facilitate back-splicing (5–8). Among these, SAM68, QKI, FUS, NF90/NF110 regulate circRNA formation through distinct mechanisms (9–11). SAM68 additionally regulates mRNA translation (12) and has been reported to act as an Internal Ribosome Entry Site (IRES) *trans*-acting factor (13,14).

At the translational level, unlike linear mRNAs, circRNAs lack both a 5′ cap and a 3′ poly(A) tail, which are central features of canonical eukaryotic translation initiation (15). Consequently, translation from circular RNA templates requires alternative mechanisms capable of recruiting the translational machinery internally. These mechanisms include internal ribosome entry sites (IRESs), N^6^-methyladenosine (m^6^A)-dependent initiation, and other forms of cap-independent ribosome recruitment (16–18). IRES elements are RNA sequences that promote translation initiation internally, independently of the 5′ cap (19). These elements were first discovered in viral RNAs, but subsequent studies have identified putative IRESs in mammalian mRNAs (20), indicating that internal initiation may also regulate cellular protein expression, especially under certain physiological or stress conditions (21–23). Notably, short IRES-like elements driving cap-independent translation are moreover enriched in endogenous circRNAs relative to linear mRNAs (24), making circRNAs a promising source of sequences capable of recruiting the translational machinery internally. However, the identification of IRESs, particularly cellular ones, remains largely empirical, as no definitive sequence or structural signature has been established (25,26).

IRES-mediated initiation does not necessarily rely on a single conserved mechanism. Depending on the IRES sequence and structural context, internal initiation can involve direct or indirect recruitment of ribosomal subunits and translation initiation factors. RBPs known as IRES trans-acting factors (ITAFs) can contribute to this process by binding IRES elements and influencing their structure and interaction with components of the translational machinery (25,27). IRESs have been studied almost exclusively as *cis*-acting elements, embedded in the same molecule as the coding region they regulate. IRES activity can, however, also be exerted *in trans*. IRESs can act as effector domains in linear and circular SINEUP RNAs, and the IRES-containing circRNA *circMyc* promotes translation of an endogenous target mRNA recognized through antisense complementarity (28). In SINEUPs (SINEB2 sequence to UP-regulate translation), a binding domain that recognizes the target mRNA through sequence complementarity is coupled to an effector domain that enhances its translation (28,29). For example, a synthetic SINEUP targeting *FXN* mRNA increased frataxin expression and rescued associated defects in a cellular model of Friedreich’s ataxia (30). The same organization is used in engineered translation-activating RNAs (taRNAs), in which an IRES-derived effector domain is fused to a sequence complementary to a selected mRNA (31). We therefore asked whether a circRNA-derived IRES-like element could be fused to an antisense sequence recognizing a selected region of a target mRNA and used to increase the expression of that target. Most RNA-based tools reduce gene expression, whereas strategies for selectively increasing protein levels without modifying the gene remain scarce. Translational activators could address this gap by enabling controlled increases in protein expression, particularly in haploinsufficient and dosage-sensitive conditions. They could also support protein homeostasis by increasing the availability of molecular chaperones. These proteins help maintain proteostasis, the coordinated processes of protein synthesis, folding, trafficking, quality control, and degradation (32,33). Disruption of this network can lead to the accumulation of misfolded or aggregation-prone proteins, a common feature of many pathological conditions, particularly neurodegenerative diseases. Heat shock proteins (HSPs) are key components of this network, making them relevant targets for translational upregulation (33,34). Their expression is modulated in response to diverse cellular stresses, including heat shock, hypoxia, oxidative stress, and ischemia, reflecting their central role in maintaining protein homeostasis under such conditions (35). Among the cytosolic chaperone systems, HSP90 has been directly implicated in protein aggregation in several neurodegenerative contexts; it inhibits α-synuclein aggregation and reduces Tau fibrillization and participates in the regulation of other aggregation-prone proteins associated with neurodegeneration (36,37). Interestingly, HSP90 inhibition has also been investigated as a therapeutic strategy in tauopathies and other proteinopathies, in part because relieving HSP90-mediated repression of HSF1 can induce a broader heat-shock response (38,39). In contrast, our approach aims to directly increase HSP90 abundance to enhance its chaperone activity toward specific aggregation-prone clients, providing a complementary strategy to approaches that modulate the heat-shock response indirectly through HSP90 inhibition.

Another example includes HSP70, that orchestrates a distinct network of co-chaperones including HSP40, to control protein misfolding and aggregates (40–42). Amyotrophic Lateral Sclerosis (ALS) is a progressive neurodegenerative disease characterized by the degeneration of motor neurons and dysregulation in RNA metabolism, stress responses, and protein homeostasis (43). Among the proteins strongly associated with ALS is fused in sarcoma (FUS), an RBP involved in transcriptional regulation, RNA processing, transport, and stress-granule biology (44–46). Under physiological conditions, FUS is predominantly localized to the nucleus. Several disease-associated mutations can disrupt its nuclear localization and promote cytoplasmic accumulation. The P525L mutation is particularly severe because it affects the nuclear localization signal (NLS), leading to defective nuclear import and cytoplasmic accumulation of FUS (47). Cytoplasmic FUS can participate in stress granule formation and undergo abnormal phase separation, creating conditions that favor the maturation of dynamic condensates into persistent protein aggregates (48,49). FUS pathology has been associated with activation of cellular stress responses and increased expression of molecular chaperones, including HSP70 and HSP90 (49). HSP90 participates in the regulation of aggregation-prone proteins in neurodegenerative diseases such as Parkinson’s and Alzheimer’s diseases (36,37), yet its effect on cytoplasmic FUS aggregation has not been tested.

In this study, we characterized a circRNA-derived IRES-like element, *SynFarsa*, as a tool for translational regulation. We first characterized its ability to promote cap-independent translation *in cis*, then fused it to antisense guide sequences, to target endogenous mRNAs through their UTRs. As a functional application, we selected HSP90 because of its central role in cellular proteostasis and its established involvement in the control of aggregation-prone proteins. We therefore investigated the effect of increased HSP90 levels on cytoplasmic FUS aggregates in a cellular model of ALS.

## Results

We aim at identifying a circRNA-derived sequence able to act as a translational effector domain. circRNAs are enriched in short IRES-like elements that support cap-independent translation (24). We focused on the SAM68-dependent circRNAs previously characterized by our group (9), since SAM68 regulates RNA processing and translation and can act as an ITAF (12–14,50). Using circAtlas (51) and IRESFinder (52), we screened 27,110 circRNAs and identified 16,355 with high-scoring predicted IRES-like elements (see Materials and Methods). Comparison with our set of 63 SAM68-regulated circRNAs yielded three candidates (**Fig. 1A**). To prioritize among the three candidates, we used *cat*RAPID (53) to predict their interactions with RBPs involved in translation, in both mouse and human (54). *mmu-Farsa_0002* showed the highest predicted propensity to interact with translation-related RBPs, and therefore with potential ITAFs, in both species (**Fig. 1B** and **S1A**). Applying the same analysis to the two circAtlas annotations of this circRNA, *mmu-Farsa_0002* (v2.0) and *mmu-Farsa_0003* (v3.0), indicated a higher propensity for the former (**Fig. 1C** and **S1B**), which we selected for experimental characterization. The corresponding 338-nucleotide sequence, generated by circularization of *Farsa* exons 5-7, is referred to hereafter as *SynFarsa*.

**Figure 1:**
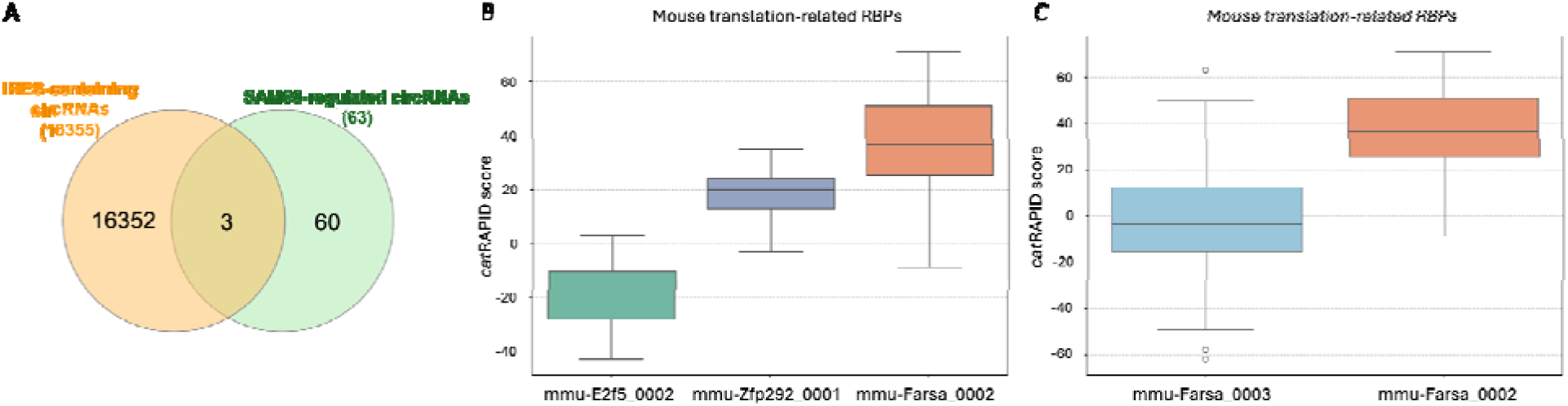
Identification of *mmu-Farsa_0002* as a circRNA-derived IRES-like element. **A**: Venn diagram combining circRNAs containing IRES elements with SAM68-regulated circRNAs. **B**: Propensity to interact with mouse translation-related RBPs of the three circRNAs candidates (*mmu-Farsa_0002, mmu-Zfp292_0001* and *mmu-E2f5_0002*). On the y axis the interaction propensity score calculated using *cat*RAPID is shown for each RBP. **C**: *mmu-Farsa_0002* (v2.0) and *mmu-Farsa_0003* (v3.0) propensities to interact with mouse translation-related RBPs. On the y axis the interaction propensity score calculated using *cat*RAPID is shown for each RBP.

### *SynFarsa* exhibits an IRES-like activity *in cis*

The putative IRES-like activity of *SynFarsa* was first evaluated using a dual-luciferase bicistronic reporter system (19,55,56). In this construct, the candidate sequence was inserted between the coding sequences of *Renilla* luciferase (*Rluc*) and *Firefly* luciferase (*Fluc*) (**Fig. 2A**). The translation of the first open reading frame (ORF), *Rluc*, occurs through cap-dependent translation, while the initiation of translation for the second ORF, *Fluc*, relies on the putative IRES inserted between the two ORFs (**Fig. 2A**). To evaluate the IRES activity of *SynFarsa*, the *cmyc* IRES (28,57) served as a positive control, and the empty plasmid (pRUF) functioned as a negative control, respectively. As an additional negative control (NegCtrl), we selected a sequence having the same length as the *SynFarsa* sequence but not predicted to be an IRES. We detected an increase in *Fluc* activity in the presence of *cmyc* and *SynFarsa* IRES without affecting *Firefly* or *Renilla* RNA levels (**Fig. 2B and S2A**), while NegCtrl did not exhibit any effect on *Fluc* translation (**Fig. 2C**). This result suggests that *SynFarsa* harbors an IRES-like activity in *cis*. We next asked whether any region of *SynFarsa* is dispensable for its IRES-like activity and tested two truncated versions of the sequence (**Fig. 2A**). The first fragment retained partial *Fluc* activity, whereas the second, lacking a larger portion of the sequence, showed no activity above the control (**Fig. 2D**), suggesting that perturbations in the sequence likely impact the structural integrity, influencing IRES function.

**Figure 2:**
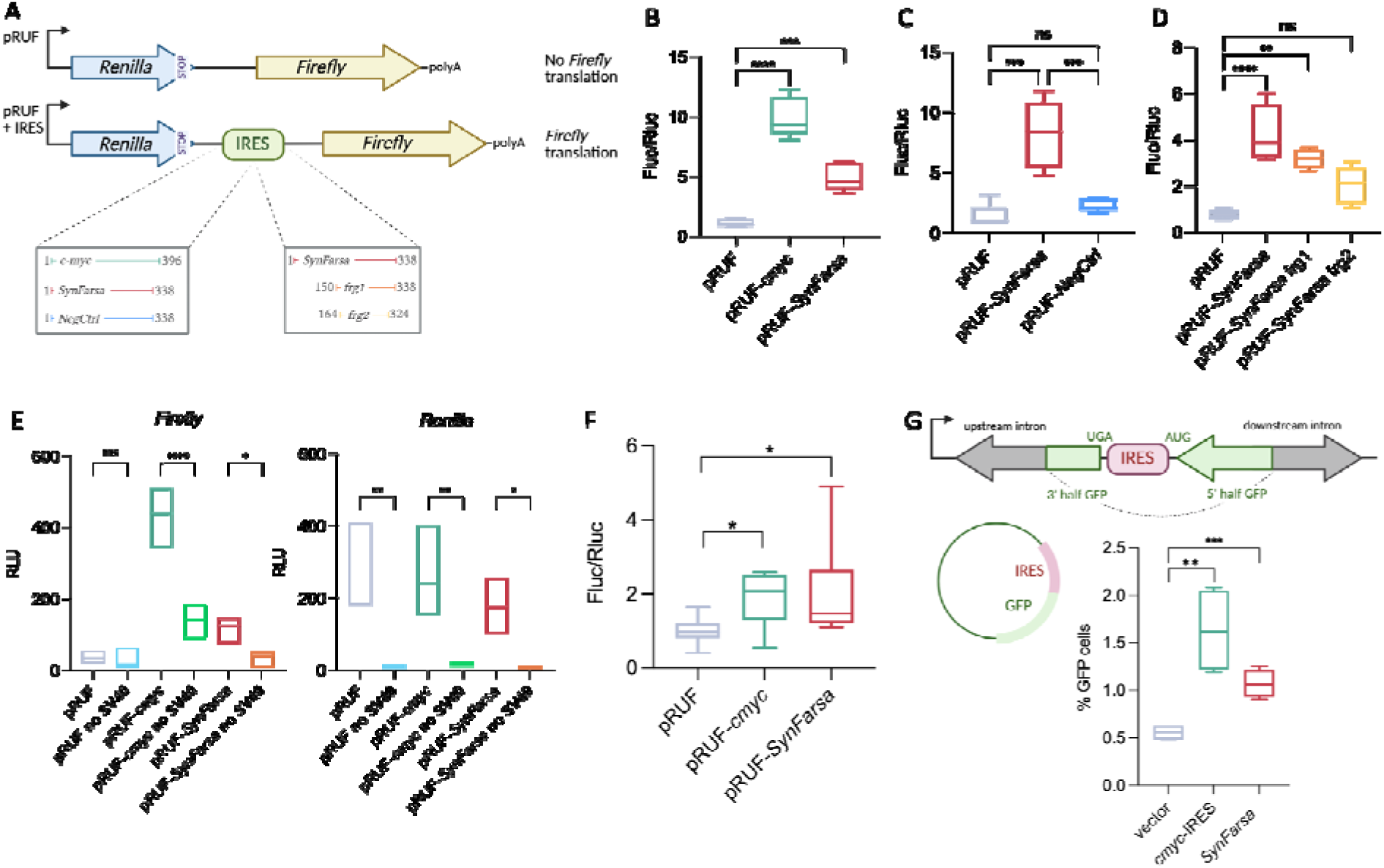
*SynFarsa* exhibits an IRES-like activity *in cis*. **A**: Schematic representation of the pRUF dual luciferase bicistronic vector showing the IRESs and fragments of interest cloned between *Renilla* and *Firefly* cistrons. *Cmyc* IRES represents the positive control, *SynFarsa* represents our IRES candidate under investigation, NegCtrl represents the negative control lacking IRES activity and *frg1* and *frg2* represent two distinct fragments of *SynFarsa*. B: *Fluc* activity normalized to *Rluc* (*Fluc/Rluc*). The impact of *SynFarsa* and *cmyc* on *Fluc* translation was assessed by normalizing the signal from each construct to that of pRUF. Box plots and lines at the median represent data from 5 biological replicates, and significance was assessed through ordinary one-way ANOVA (p<0.05). **C**: *SynFarsa* activity compared to the negative control NegCtrl. Box plots and lines at the median represent data from 5 biological replicates, and significance was assessed through ordinary one-way ANOVA (p<0.05). **D**: fragmentations of the *SynFarsa* sequence reduce its IRES-like activity. The impact of each fragment on *Fluc* translation was assessed by normalizing the signal to that of pRUF. Box plots and lines at the median represent data from 5 biological replicates, and significance was assessed through ordinary one-way ANOVA (p<0.05). **E**: Relative light units (RLU) for *Firefly* and *Renilla* luminescence, following transfection of pRUF plasmids with or without the SV40 promoter. The luminescence signal is notably reduced in the absence of the SV40 promoter. In the case of *Fluc* signal, there is no significant difference between pRUF and pRUF without SV40, as no IRES sequence is present to support *Fluc* translation. Box plots and lines at the median represent data from 5 biological replicates, with significance determined via one-way ANOVA (p<0.05). **F**: *Fluc* activity normalized to *Rluc* (*Fluc/Rluc*) and measured upon transfection of bicistronic RNAs *in vitro* transcribed. The functionality of *cmyc* and *SynFarsa* IRESs was assessed by normalizing the signal to the pRUF RNA control. Box plots and lines at the median represent data from 8 biological replicates. Significance was determined using unpaired t-tests (p<0.05). **G**: schematic illustration of the circularizing GFP reporter system (top). *circGFP* is translated only when the sequence inserted between the separated GFP fragments functions as an IRES. The percentage of GFP-positive cells as determined by flow cytometry. Box plots and lines at the median represent data from 4 biological replicates, with significance determined via unpaired t-test (p<0.05) (bottom).

Different artefacts may contribute to the false discovery of IRES sequences, for instance, the presence of a cryptic promoter within the IRES under investigation that initiates alternative transcription, producing an mRNA comprising only the second ORF or unpredicted splicing events that generate in-frame fusion ORFs (19,55,58). To address these potential artefacts, we carried out several additional controls. First, we conducted a dual-luciferase reporter plasmid eliminating the SV40 promoter from the pRUF vector. Deletion of the SV40 promoter resulted in a significant decrease in both *Rluc* and *Fluc* activity (**Fig. 2E**), demonstrating that the IRES activity *in cis* is not a consequence of the *SynFarsa* IRES sequence acting as a cryptic promoter. Second, the *in vitro* bicistronic transcripts were transfected into cells. This bypasses transcription from the plasmid and therefore excludes both cryptic promoter activity and splicing events that could generate an independent *Fluc* transcript carrying its own 5′ end, which would be translated in a cap-dependent manner and could thus account for the *Fluc* signal. Under these conditions, both the *cmyc* IRES and *SynFarsa* induced *Fluc* translation (**Fig. 2F**), while *Renilla* and *Firefly* RNA levels were comparable across conditions (**Fig. S2B and S2C**). The reduced activity in this assay compared to the vector transfection experiment could be attributed to the fact that these IRES-containing RNAs are not synthesized in the nucleus, and they may lack association with crucial factors (*e.g.* ITAFs) for IRES activity (59). Lastly, we used a circRNA expressing GFP to measure IRES activity (**Fig. 2G**). The GFP ORF is divided into two fragments in reverse order, ensuring that the intact ORF can only be reconstituted upon the formation of the circRNA through back-splicing. Expression of GFP can occur only if the sequence cloned between the two GFP fragments acts as an IRES. Both *cmyc* IRES and *SynFarsa* were both able to sustain *circGFP* translation in comparison to the negative control, *i.e.* plasmid lacking the putative IRES sequence (pRUF) (**Fig. 2G**). GFP protein expression was detected by Western blot (**Fig. S2D**) and circular and linear RNA levels were measured by RT-qPCR analysis (**Fig. S2E**). We concluded that *SynFarsa* harbors an IRES-like activity capable of activating translation in *cis*.

### *SynFarsa* targeting the 3′UTR of *hsp90* increases HSP90 protein expression

Having established that *SynFarsa* exhibits IRES-like activity *in cis*, we next asked whether this activity could be repurposed as a programmable system for modulating the expression of endogenous proteins, using a design based on the SINEUP architecture (28). The design principle was to combine the *SynFarsa* effector domain, which recruits the translational machinery, with a sequence-specific RNA guide (gRNA) directed against a selected region of the target mRNA (**Fig. 3A**) (28,60). We first confirmed that *SynFarsa* induces *Fluc* translation in human cells (**Fig. S3A**). As a proof of concept, we selected *hsp70* and *hsp90* mRNAs as target transcripts. To enable sequence-specific recognition, regions within the 5′ and 3′ UTRs of *hsp70* and *hsp90* transcripts were analyzed using RNA secondary-structure prediction. Candidate regions with relatively low structural complexity were preferentially selected, avoiding highly structured double-stranded regions. Because mammalian 5′ UTRs are shorter than 3′ UTRs (61), 13-nucleotide fragments were selected from the 5′UTRs, whereas 30-nucleotide fragments were selected from the 3′ UTRs. For each target region, the reverse-complementary sequence was generated and fused downstream of the *SynFarsa* IRES-like element, thereby creating the *SynFarsa*-gRNA molecule. Fusing the gRNA sequences to S*ynFarsa* did not compromise its IRES-like activity (**Fig. S3B and S3C**). For constructs targeting both UTRs simultaneously, a 40-nucleotide linker was introduced between the targeting regions to provide sufficient stability while minimizing the formation of complex secondary structures.

**Figure 3:**
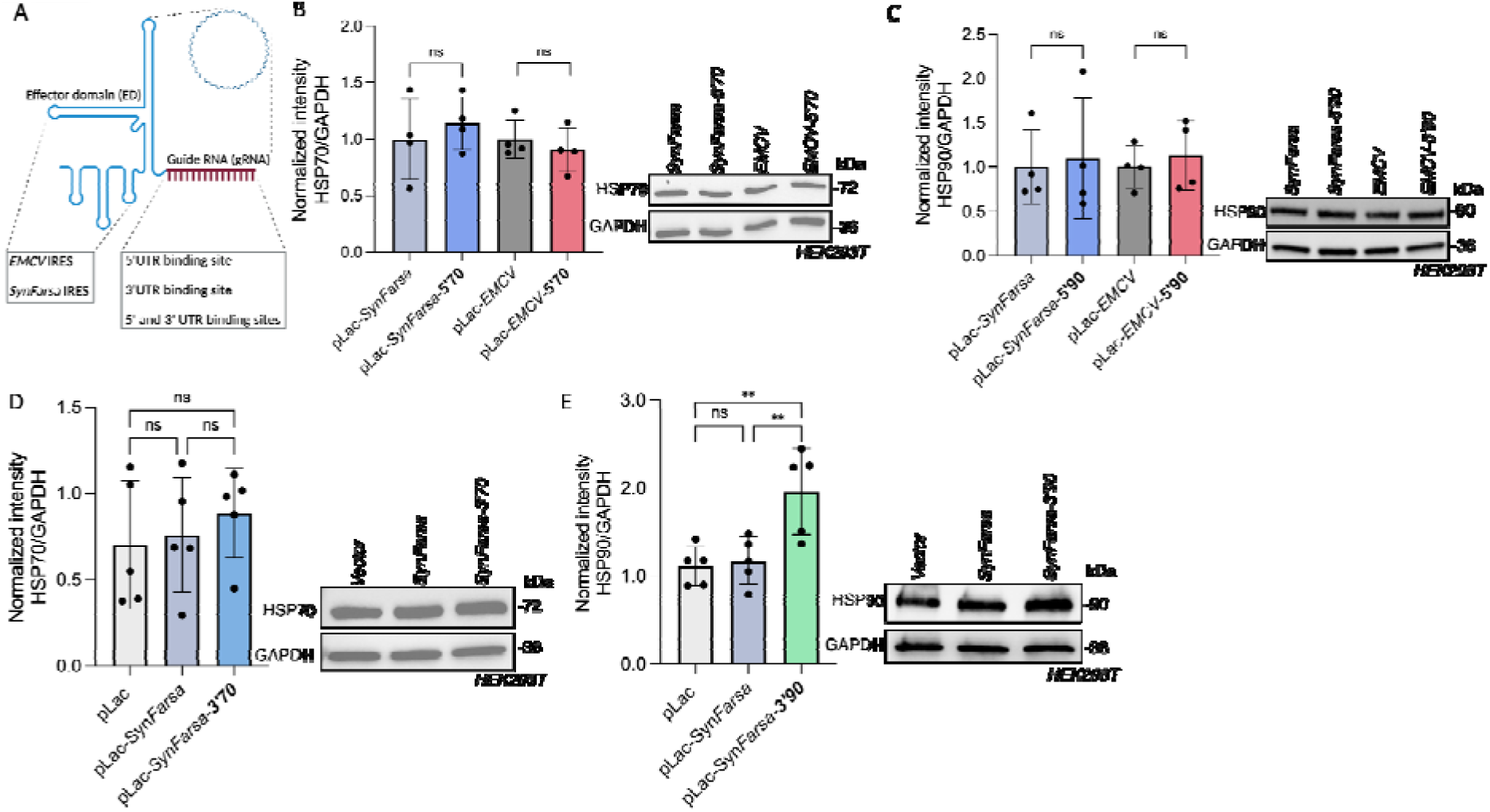
The *SynFarsa* model for targeted translational activation of endogenous mRNAs. **A**: schematic representation of the *SynFarsa* circular model. **B** and **C**: Representative western blots showing that targeting the 5’UTR of *hsp70* and *hsp90* did not change HSP70 (**B**) and HSP90 (**C**) protein levels in HEK293T cells following 48 hours of transfection. Quantifications normalized to the loading control GAPDH are shown as plots on the left, and representative blots are shown on the right. N= 4 biological replicates and non-significance was assessed through t-test (p<0.05). **D** and **E**: Representative Western blots showing that targeting the 3’UTR of *hsp70* did not change HSP70 protein levels (**D**), whereas targeting the 3’UTR of *hsp90* resulted in a ∼1.6-fold increase in HSP90 protein expression (**E**). Quantifications normalized to the loading control GAPDH are shown as plots on the left, and representative blots are shown on the right. N= 5 biological replicates and non-significance was assessed through t-test (p<0.05) (**D**). N= 5 biological replicates and significance was assessed through t-test (p<0.05) (**E**).

We first investigated whether the engineered *SynFarsa* could enhance the expression of endogenous HSP proteins. The first strategy consisted of targeting the 5’UTR of *hsp70* and *hsp90* mRNAs, where *SynFarsa* and the *EMCV* positive control were used as effector domains. Protein levels were assessed by Western blotting after 48 hours of transfection and normalized to GAPDH. Targeting the 5′UTRs of either *hsp70* or *hsp90* with *SynFarsa* did not significantly alter protein expression (**Fig. 3B** and **3C**) or the corresponding mRNA levels (**Fig. S3D** and **S3E**). The *EMCV* IRES used as a positive effector control was also inactive at these sites, suggesting that these targeting configurations may not support translational activation. We next redirected *SynFarsa* to the 3′UTRs of *hsp70* and *hsp90* mRNAs. Targeting the *hsp70* 3′UTR had no detectable effect on HSP70 protein levels (**Fig. 3D**), whereas targeting the *hsp90* 3′UTR increased HSP90 expression (**Fig. 3E**). The *hsp70*-targeting constructs provide an internal control: they carry guides of the same length, at the same position within the same backbone, fully complementary to an endogenous transcript, yet did not increase HSP70 levels. This suggests that translational activation depends on the target context rather than simply on the addition of a complementary guide.

To test whether this increase reflected a higher abundance of the target transcript, we measured endogenous *hsp90* mRNA by RT-qPCR and detected no significant change (**Fig. S3F**). Having identified the *hsp90* 3′UTR as an effective target region, we next asked whether simultaneous recognition of the 5′ and 3′UTRs could further enhance *SynFarsa*-mediated translation. Despite carrying two target-recognition sequences, this construct did not increase HSP90 protein levels relative to the control condition (**Fig. S3G**). Finally, resistance to RNase R digestion indicated that *SynFarsa*-3′90 undergoes circularisation (**Fig. S3H**). This assay does not, however, establish the relative abundance of the circular and linear species, nor which form mediates translational activation.

### Upregulated HSP90 reduces FUS aggregates

Having established that *SynFarsa-3*′*90* can increase endogenous HSP90 expression, we next investigated whether this targeted upregulation of a molecular chaperone could influence protein aggregation. To address this question, we used a neuroblastoma cellular model expressing the ALS-associated FUS^P525L^ mutant (45) (**Fig. S4A**). Following doxycycline induction, cells were exposed to a 1hour treatment with sodium arsenite (SA) to induce cellular stress and promote the formation of cytoplasmic aggregates. Because molecular chaperones are central components of the cellular response to proteotoxic stress (62), we first examined whether HSP90 expression was altered under the aggregation-inducing conditions. Aligning with previous observations in the FUS^P525L^ model (49), SA treatment resulted in a slight but statistically significant increase in HSP90 protein levels, as determined by Western blot analysis (**Fig. 4A**). This increase was further supported by immunofluorescence analysis, which showed an increase in HSP90 fluorescence following stress induction (**Fig. 4B and S4B**). We therefore reasoned that further increasing HSP90 expression using the *SynFarsa-3*′*90* translational system could enhance this endogenous response and potentially influence FUS aggregation. To do that, we transfected FUS^P525L^-expressing SK-N-BE cells with *SynFarsa-3*′*90*, using *SynFarsa* as control, and exposed the cells to SA. Western blot analysis showed that *SynFarsa-3*′*90* increased endogenous HSP90 protein levels by approximately 1.4-fold compared with the *SynFarsa* control in stressed cells (**Fig. 4C**). The same upregulation was observed in cells that had not been exposed to SA, as in the earlier experiments (**Fig. 3E**). *SynFarsa*-3′90 therefore increases HSP90 levels independently of stress induction. These results indicate that the targeted translational activity of *SynFarsa-3*′*90* is retained in the FUS^P525L^ neuroblastoma model and under the stress conditions used to induce protein aggregation. Having confirmed that *SynFarsa-3*′*90* enhances HSP90 expression in the FUS^P525L^ model, we next asked whether this increase was accompanied by a change in FUS aggregation. Quantification of confocal immunofluorescence images showed that cells expressing *SynFarsa*-3′90 contained fewer cytoplasmic FUS aggregates than control cells expressing *SynFarsa* (**Fig. 4D and 4E**). Cytoplasmic FUS aggregates were also smaller in cells expressing *SynFarsa-3*′*90* than in the *SynFarsa* control (**Fig. S4C**). Together, these results show that *SynFarsa* can upregulate HSP90 expression and reduce cytoplasmic FUS aggregation.

**Figure 4:**
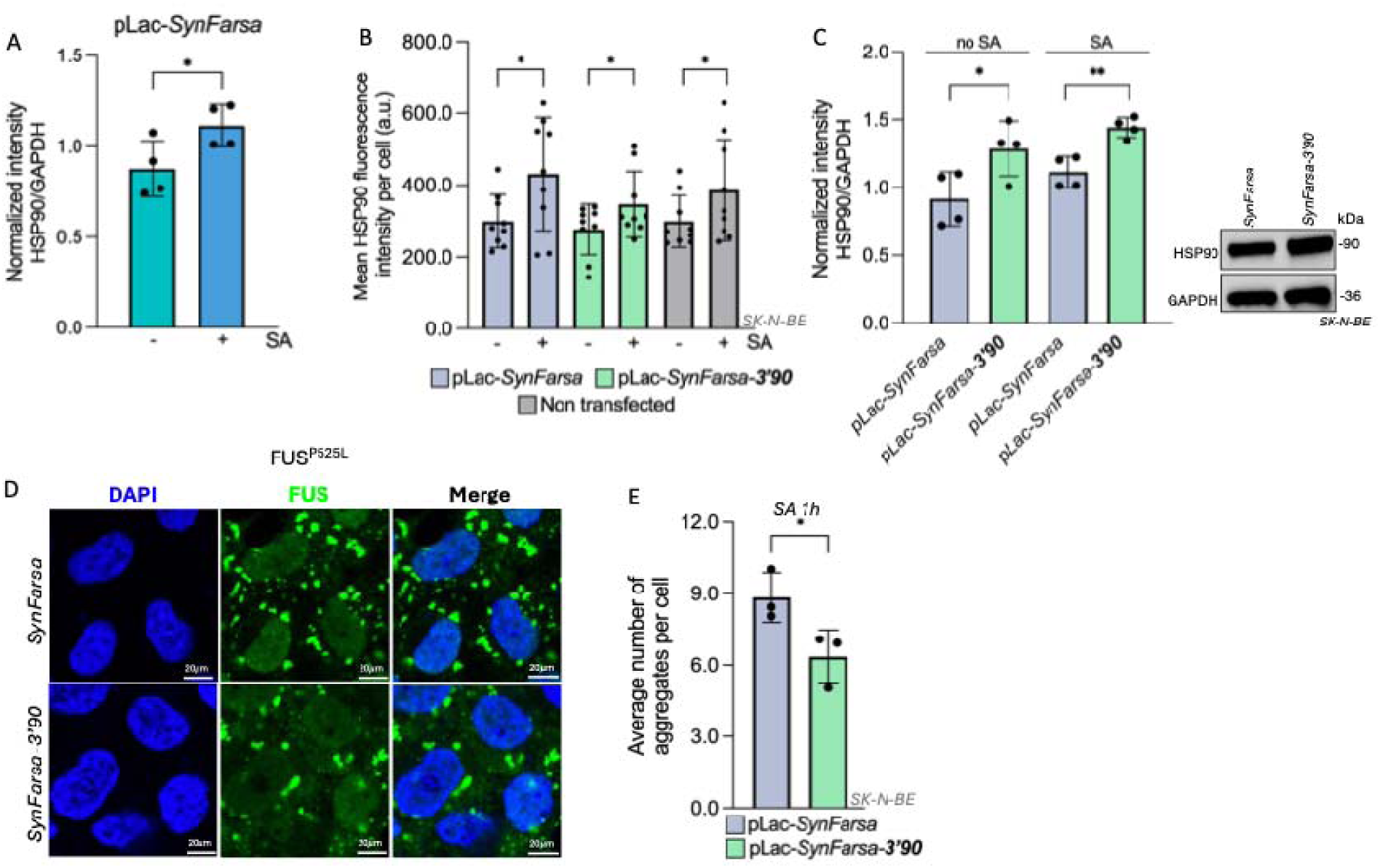
*SynFarsa*-mediated HSP90 upregulation modulates FUS aggregation. **A**: representative Western blots showing upregulation of HSP90 protein levels after SA stress induction. Quantification normalized to the loading control GAPDH is shown as bar plots for N= 4 biological replicates. Significance was assessed through unpaired t-test (p<0.05). **B**: HSP90 fluorescence intensity quantified using a specific FIJI-JAVA script. The intensity is quantified using DAPI-defined nuclear ROIs. Mean fluorescence intensity was measured for each cell, and values were averaged per image for statistical comparison between conditions. Significance was assessed using unpaired t-test (p<0.05) for technical images deriving from 3 independent biological replicates. **C**: Western blot analysis confirming HSP90 upregulation upon *SynFarsa*-3’90 transfection, in stressed and non-stressed conditions. Quantification bar plots normalized to the loading control GAPDH are on the left and representative blots on the right. N= 4 biological replicates and significance was assessed through unpaired t-test (p<0.05). **D**: representative confocal images of FUS^P525L^-expressing SK-N-BE cells. DAPI staining, FUS^P525L^ expression and merged conditions subjected to 1 hour of SA stress are shown. Scale bar = 20 μm. **E**: average number of total aggregates quantification. Bar plots represent values from 3 independent biological replicates.

## Discussion

Here we developed a synthetic circular RNA tool that harnesses the IRES-like activity of *SynFarsa* to modulate endogenous protein expression. Identifying such elements remains a challenge as no definitive sequence or structural signature defines an IRES (54), making experimental validation essential. circRNAs offer a promising starting point, short IRES-like elements driving cap-independent translation are enriched in endogenous circRNAs relative to linear mRNAs, and most circRNAs are predicted to contain at least one (24). Together with extensive circRNA catalogues (51), these observations provide a basis for exploring circRNAs as a source of translational effector sequences. An important consideration is whether this activity is strand-specific. In translational reporter assays, inversion of a candidate sequence is commonly used to test whether activity requires the element itself (63). However, the reverse complement of a structured RNA is also expected to be structured (64), and our analysis identified predicted IRES-like features in both orientations for a substantial proportion of candidates. Strand specificity should therefore be verified experimentally when evaluating cellular or circRNA-derived IRES-like elements. Demonstrating IRES-like activity requires distinguishing genuine internal translation initiation from reporter artefacts. Bicistronic and back-splicing systems can produce false-positive signals through promoter activity within the candidate sequence or aberrant RNA processing. In particular, split-GFP back-splicing reporters can generate linear *trans*-spliced transcripts that account for a substantial fraction of the detected reporter protein (19,56,65,66). We therefore tested *SynFarsa* using three complementary approaches. Deleting the SV40 promoter reduced *Firefly* activity from the *SynFarsa*-containing construct, arguing against autonomous transcription of the downstream cistron. Crucially, *SynFarsa*-dependent activity persisted after transfection of *in vitro*-transcribed bicistronic RNA, bypassing both plasmid-borne promoters and splicing. The circular GFP reporter supported these findings, although a contribution from linear RNA species cannot be excluded in this assay (65,67).

We next asked whether this activity could be exploited *in trans* and converted *SynFarsa* into a regulator directed against a selected endogenous transcript. *SynFarsa-3*′*90* combines a binding domain conferring target specificity with an effector domain promoting translation, following the architecture of a circular SINEUP (28). The absence of free ends is expected to confer greater resistance to exonucleases and improve RNA stability, while the circular configuration may also influence the folding and activity of the circRNA-derived effector. However, the contribution of circularity to stability and translational activation remains to be determined by comparison with a linear counterpart.

Our observation builds on recent evidence that IRESs can function as effector domains in linear and circular SINEUPs (28), as well as in engineered translation-activating RNAs directed against endogenous transcripts (31). How these molecules stimulate translation *in trans* remains an open question. Reported mechanisms include increased association of the target mRNA with polysomes (28) and recruitment of eIF3 and eIF4G by taRNA engineered activators (31), while the role of ITAFs remains unexplored in these systems. The predicted interactions of *SynFarsa* with translation-related RBPs suggest a possible contribution from ITAFs. Given our focus on SAM68-regulated circRNAs, SAM68 is a priority candidate to test, and its depletion would directly assess whether *SynFarsa*-mediated translational activation depends on this factor. Only one of the five configurations tested, targeting the *hsp90* 3′UTR alone, supported upregulation. Targeting the *hsp70* 3′UTR, the 5′UTR of either transcripts, or both UTRs of *hsp90* simultaneously was ineffective. The activity observed upon *hsp90* 3′UTR targeting is consistent with findings for engineered taRNAs, for which 3′UTRs were generally the most permissive regions and offered greater flexibility for guide design (31). However, our results show that targeting a 3′UTR does not guarantee activity, and adding a second binding site does not necessarily improve it. Local RNA structure, accessibility and the positioning of the effector relative to the translation machinery may contribute to these differences. Combined targeting could also alter the geometry or accessibility of the RNA complex, although this remains to be tested.

Finally, the magnitude of upregulation we obtained (1.4-fold to 1.6-fold) is comparable to that reported for other *trans*-acting translational activators (68). Such modest increases may be useful in dosage-sensitive and haploinsufficient conditions, where restoring protein levels requires controlled modulation (28,31). In the specific FUS^P525L^ neuroblastoma model, this increase in HSP90 was accompanied by a reduction in the number and size of cytoplasmic FUS aggregates, supporting the activity of the system in a disease-relevant cellular context. Although the mechanism underlying this effect remains to be established. Increased HSP90 levels could influence FUS proteostasis directly, through chaperone activity on FUS itself or through co-chaperone complexes, or indirectly, by altering the dynamics of the aggregates. However, effects unrelated to HSP90 are also possible; *SynFarsa* expression or circRNA transfection could affect aggregates formation or activate cellular RNA-sensing pathways independently of HSP90. Thus, the observed reduction in FUS aggregation cannot be attributed to HSP90 upregulation alone at this stage. Demonstrating that HSP90 inhibition or knockdown abolishes the effect of *SynFarsa* on FUS aggregation would provide direct evidence for a causal role of HSP90 in this phenotype. *SynFarsa* thus provides a circRNA-derived effector domain that, when coupled to a target-binding sequence, increases endogenous protein expression without altering mRNA abundance, following the design of an artificial circular SINEUP. Direct engagement with the *hsp90* transcript, changes in its ribosomal loading, the contribution of ITAFs and the requirement for the circular configuration remain to be established. Specificity beyond the intended target, including potential off-target hybridization, has also not been assessed. Moving towards therapeutic applications will require production and testing of *in vitro*-transcribed circRNA, an effective delivery strategy and validation in neurons or patient-derived cells. With these steps, targeted translational regulation could offer a route to restoring protein homeostasis in neurodegenerative diseases.

## Material and Methods

### Cell culture

Mouse neuroblasts Neuro-2a (N2a) and Human Embryonic Kidney (HEK) 293T cells were cultured in complete DMEM (41965039, Gibco) supplemented with 10% FBS (A5256701, Gibco), 1% Penicillin/Streptomycin (15140122, Gibco) and 1% Sodium Pyruvate (1136007, Gibco) when necessary. The SK-N-BE FUS^P525L^ cell line was provided by Bozzoni’s laboratory. Cells were cultured in complete RPMI medium (31870025, Gibco) supplemented with 10% FBS-Tetracycline free (VWR, S181T-500), 1% Penicillin/Streptomycin (15140122, Gibco), 1% L-Glutamine (25040081, Gibco) and 1% Sodium Pyruvate (1136007, Gibco). All cell lines used were tested for mycoplasma contamination and grown at 37°C, 5% CO2.

### Plasmid constructs

PCR amplification reactions were performed using CloneAmp^TM^ HiFi PCR Premix (639298, Takara Bio) or Q5 Hot Start High Fidelity 2X Mix (M0494S, New England Biolabs) following each manufacturer’s instructions. Plasmids used in this study are listed in **Supplementary Table 1**. The IRES activity was assessed through a luminescence assay using the pRUF plasmid. Both the empty pRUF and the pRUF-*cmyc* vectors were provided by Prof. Stefano Gustincich’s laboratory. For the following cloning, all the primers used are listed in **Supplementary Table 2**. To generate the pRUF plasmids that contain *SynFarsa* and fragments *Frg1* and *Frg2*, the *SynFarsa* sequence (**Supplementary Table 3**) was amplified from N2a cDNA by PCR.

The 338-nucleotides negative control, having the same length as *SynFarsa* and not predicted to have IRES activity, is derived from the 3’UTR of *TARDBP* and was amplified from a plasmid already available in our laboratory. After amplification, the amplicon was digested with NdeI and XhoI restriction enzymes, pRUF was linearized using the same enzymes, and the amplicon was cloned into linear pRUF between *Rluc* and *Fluc* using Quick Ligation^TM^ Kit.

For the promoter-less control, SV40 promoter was removed from pRUF vectors by PCR amplification using the divergent primers listed in **Supplementary Table 2**. The amplicons were purified from agarose gel using Monarch^R^ DNA Gel Extraction Kit (New England Biolabs) and then digested with Dpn1 (New England Biolabs). Linearized plasmids were then circularized using T4 Polynucleotide Kinase (New England Biolab), followed by incubation with DNA ligase Quick Ligation^TM^ Kit (New England Biolabs).

The overexpression of *SynFarsa* was assessed using the pCDNA3.1(+) LACCASE2 MCS Exon Vector, referred to by pLac. *SynFarsa* sequence was cloned in pLac following Gibson Assembly strategy from NEBuilder^R^ HiFi DNA Assembly Cloning Kit (New England Biolabs). The plasmid was linearized with divergent primers and treated with Dpn1. *SynFarsa* was amplified from cDNA obtained from reverse transcription of total N2a RNA, using primers that overlap a region of 20 nucleotides with the linearized plasmid.

The UTRs fragments of *hsp70* and *hsp90* were selected from Vienna RNAfold server, taking into consideration low-complexity regions, and their reverse complement sequences gRNA were used to design the primers for the cloning (**Supplementary Tables 3 & 4**). Primers were designed to harbor the specific reverse complement UTR fragments that were cloned by mutagenesis into pRUF and pLac vectors. For the IRES-UTR activity, luminescence assay was performed using pRUF vector where the gRNA targeting 5’ and 3’UTRs were fused to *SynFarsa* and cloned between *Rluc* and *Fluc*. To study the potential effect of *SynFarsa* on *hsp* mRNA translation, the gRNAs targeting 5’UTR, 3’UTR and both UTRs (referred to by 5’3’UTR) were cloned in pLac vector fused to *SynFarsa*. In the case of both UTRs cloning, a linker sequence (**Supplementary Table 4**) was introduced between the UTRs to prevent them from fusing during circularization.

PCR amplification of the 5’UTRs targeting gRNA was performed using CloneAmp^TM^ HiFi PCR Premix (639298, Takara Bio), following the manufacturer’s instructions. The 3’UTR and 5’3’UTR targeting gRNAs were obtained by GenScript cloned in the pUC57 plasmid (**Supplementary Table 1**) in the presence of *SynFarsa*. To isolate the fragments and clone them onto pLac vector, pLac was first linearized using divergent primers, digested with Dpn1 (New England Biolabs) and the *SynFarsa*-UTR fragments were PCR amplified from pUC57 using corresponding primers and the Q5 Hot Start High Fidelity 2X Mix (M0494S, New England Biolabs) following manufacturer’s instructions. The PCR fragments were then cloned in pLac following Gibson Assembly strategy from NEBuilder^R^ HiFi DNA Assembly Cloning Kit (New England Biolabs). To generate the constructs for the circular GFP reporter assay, the vector pcDNA3.1(+) ZKSCAN1 MCS-WT Split GFP was linearized by PCR using the divergent primers listed in **Supplementary Table 2**. The primers were designed to linearize the plasmid and, at the same time, remove the sense EMCV IRES contained in the vector. *SynFarsa* was amplified using the respective pRUF plasmids as templates with primers containing an overhang region (20 nt) overlapping to the linearized pcDNA3.1(+) ZKSCAN1 MCS-WT Split GFP at the desired site of insertion. The linearized plasmids and each of the amplicons were assembled using the NEBuilder® HiFi DNA Assembly Cloning Kit (New England Biolabs) according to the manufacturer’s instructions. To generate the vector without any IRES sequences, the linearized plasmid pcDNA3.1(+) ZKSCAN1 MCS-WT Split GFP was circularized using T4 Polynucleotide Kinase (New England Biolabs) treatment, followed by incubation with DNA ligase (Quick Ligation™ Kit; New England Biolabs). The plasmid containing the *c-myc* IRES was obtained from the lab of Prof. Stefano Gustincich.

### Luminescence assay

N2a and HEK293T cells were transfected with pRUF vectors or Bicistronic *in vitro* transcribed mRNAs and harvested 24 hours post-transfection. Cells were first washed with PBS, and luminescence was measured using the Dual-Glo® Luciferase Assay System (Promega) following the manufacturer’s instructions in a microplate reader INFINITE 200 PRO (TECAN). The empty pRUF (No IRES between *Rluc* and *Fluc*) luminescence level was used to normalize the transfection efficiency or background effects. IRES activity was calculated as the ratio of *Fluc* activity to *Rluc*. RNA was purified from harvested cells and used for RT-qPCR (**Supplementary Table 6**). All experiments were conducted with a minimum of four independent biological replicates, each including technical duplicates. The obtained values were plotted using GraphPad Prism.

### Circular GFP reporter assay

N2a cells were transfected with pcDNA3.1(+) ZKSCAN1 MCS-WT Split GFP + IRES as previously described. After PBS washing, one-fourth of the cells were used for protein and RNA extraction, while the entire remaining fraction (3/4) was designated for flow cytometry. For the latter, cells were pelleted, resuspended in 1 mL of PBS supplemented with 2% heat-inactivated FBS (Gibco), and 1 mM EDTA pH 8 (ThermoFisher Scientific). Subsequently, 750 μl were filtered using Falcon filter-top tubes (ThermoFisher Scientific) to prevent clumping. Flow cytometry was performed on a Sony SH800 CellSorter; the percentage of GFP-positive cells was determined using a minimum of 200000 events recorded. The obtained values were plotted using GraphPad Prism.

### *In vitro* transcribed RNA transfection

pRUF bicistronic vectors of interest were linearized with Not I enzyme and DNA was extracted (using Monarch^®^ DNA Gel Extraction Kit), dephosphorylated (with Quick CIP NEB #M0525) and purified (using Monarch^®^ PCR and DNA Clean Up Kit). *In vitro* transcription of bicistronic mRNA was performed using mMESSAGE mMACHINE^TM^ T7 ULTRA Transcription Kit (Thermo Fisher Scientific, AM1345). *In vitro* transcribed RNA was purified using the RNeasy Mini Kit (Qiagen, 74104). N2a cells were plated one day before transfection and then transfected with the RNA following the Lipofectamine™ MessengerMAX™ Reagent protocol. Cells were harvested 6 hours after transfection by resuspending them in 100 µl of PBS and collecting in a manner of 1/3 for luminescence analysis and 2/3 for RNA analysis (**Supplementary Table 6**). Experiments were performed in the same way for at least 8 independent biological replicates. The obtained values were plotted using GraphPad Prism.

### Protein extraction and Western blot

HEK2937T cells were plated in a 6-well plate at a final concentration of 12.5 × 10^4^ per well in 2 mL of complete DMEM. 2000 ng of plasmid were transfected using Lipofectamine^TM^ 2000 transfection reagent (ThermoFisher Scientific) following the manufacturer’s protocol, and cells were kept in optiMEM for 48 hours.

SK-N-BE cells were plated in a 6-well plate at a final concentration of 12.5 × 10^4^ per well in 2 mL of RPMI medium. 2000 ng of plasmid were transfected using Lipofectamine^TM^ 2000 transfection reagent (ThermoFisher Scientific) following the manufacturer’s protocol, and cells were kept in medium without antibiotics. 4 µl of DOX (50 µg/mL) was directly added to the DOX conditions and cells were incubated for 24 hours. Before collection, stress was applied by adding 2 µl of sodium arsenite (0.5 M) and incubated at 37°C for 1 hour.

Proteins from HEK293T and SK-N-BE cell lines were extracted by resuspending pellets in RIPA buffer (25 mM Tris-HCl pH 8, 1 mM compEDTA, 1% Triton-X 100, 150 mM NaCl, 0.1% SDS and 1 mM DTT) and kept on ice for 15 minutes. Cells were centrifuged at 13000 rpm at 4°C for 10 minutes. The supernatant was preserved, and protein concentration was measured using BCA assay following the manufacturer’s instructions. 15 μg of protein from HEK293T and 5 μg from SK-N-B were mixed with LB 6X and boiled for 5 minutes before being loaded on gel. Proteins were loaded onto 4-15% Mini-PROTEAN^R^ TGX^TM^ precast protein gel (4561083, Biorad) immersed in Tris-Glycine SDS running buffer (1610732 Biorad), run at 220 V and transferred on a nitrocellulose membrane (Bio-Rad). The membrane was then blocked with 5% milk in TBS-T (TBS buffer with 0.1% Tween20) for 1 hour at RT. Primary antibody (**Supplementary Table 5**) incubation was performed in 5% milk TBS-t for 1 hour at RT, the membranes were next washed 3 times with TBS-T and incubated with secondary antibody (**Supplementary Table 5**) for 1 hour RT. To reveal the signal, the membrane was incubated with Immobilon^®^ ECL UltraPlus Western HRP Substrate (WBULP, Merck) and revealed on a ChemiDoc MP imaging system (Bio-Rad). Signal was quantified using Fiji (69) and normalized to GAPDH. The obtained results after quantification were plotted using GraphPad Prism.

### RNA extraction and RT-qPCR analysis

Total RNA from N2a and HEK293T cells was extracted using Direct-zol RNA Miniprep Kit (Zymo Research, R2052) according to the manufacturer’s instructions with 30 minutes of on-column DNAse digestion. 320 ng of RNA were reverse transcribed using PrimeScript RT Reagent Kit (Takara Bio). qRT-PCR was performed using PowerUp SYBR Green Master Mix reagent (ThermoFisher Scientific) according to the manufacturer’s instructions and QuantStudio 5 real-time PCR system with a 384-well block. RNA levels are relative to *Gapdh* mRNA and RNA quantity was calculated as the fold change (2^-ΔΔCt^) with respect to the average of the control sample. A complete list of the oligonucleotides used for RT-qPCR experiments is provided in **Supplementary Table 6**. The obtained values were plotted using GraphPad Prism.

### RNAse R treatment

Total RNA was extracted as mentioned before and 1.5 µg were treated with 10 U RNase R (Tema Ricerca, RNR07250) in 10 µL reactions at 37°C for 20 minutes, with untreated RNA serving as a control. From both treated and untreated RNA, 320 ng was reverse transcribed using the PrimeScript retro transcription master mix (Takara, RR036A). RNA abundance was quantified by qRT-PCR using 10 ng of cDNA per well for QuantStudio 5 real-time PCR system with a 384-well block (**Supplementary Table 6**). The percentage of RNA remaining after treatment was calculated using the ΔCt method according to the formula 2^(Ct untreated−Ct RNase R) x 100. The obtained values were plotted using GraphPad Prism.

### Immunofluorescence and confocal microscopy imaging

SK-N-BE FUS^P525L^ cells were plated on glass coverslips in a 24-well plate, coated with attachment factor (Merck, 123-100), at a final concentration of 1 × 10^5^ cells per well in 500 µl of RPMI medium without antibiotics. 160 ng of plasmid were transfected using Lipofectamine^TM^ 2000 transfection reagent (ThermoFisher Scientific) following the manufacturer’s protocol, and cells were kept in medium without antibiotics. 1 µl of DOX (50 µg/mL) was directly added to the DOX conditions and cells were incubated for 24 hours. Before collection, cells were subjected to oxidative stress by treatment with sodium arsenite (0.5 M) for 1 hour at 37°C, then washed, fixed in 4% paraformaldehyde for 10 minutes at RT and washed again 3 times in complete PBS. Permeabilization was performed by incubating the coverslips in PBS supplemented with 0.1% Triton X-100 for 10 minutes at RT. Samples were then blocked in PBS supplemented with 1% bovine serum albumin (BSA) for 20 minutes at RT. Next, anti-HSP90 (1:100) and anti-FLAG (1:500) primary antibodies (**Supplementary Table 5**) were diluted in 1% BSA and incubated with the samples for 1 hour at RT. Cells were then washed 3 times with PBS before being incubated with fluorophore-conjugated secondary antibodies (**Supplementary Table 5**) diluted 1:500 in 1% BSA and incubated for 1 hour at RT. After 3 PBS washes, nuclei were stained with DAPI (Merck, D9542-1MG) diluted 1 μg/mL in PBS for 5 minutes at RT and coverslips were mounted with Prolong Diamond Mounting Media (Thermo Fisher, P36965) on microscope slides.

Fixed samples were imaged using a Nikon Instrument A1MP R-S confocal laser microscope equipped with 60x objectives. Confocal images were collected with NIS-element AR software (Nikon) and analyzed with FIJI-ImageJ using a specific automated batch-processing JAVA pipeline. For HSP90 fluorescence quantification, nuclei were identified by DAPI staining and used to define the corresponding cellular ROIs. HSP90 fluorescence intensity was quantified within the corresponding ROIs in the HSP90 channel, and the mean intensity per cell was calculated. Image analysis was done using the AggreCount plugin open-source software (70). Aggregates were identified based on predefined size (0.2-20 µm²). The obtained values after quantification were plotted using GraphPad Prism.

### Statistical analysis

Statistical analyses were performed using GraphPad Prism. Biological replicates were defined as independent experiments performed on separate days, including independent cell transfections; technical replicates and individual images from the same experiment were not considered independent biological replicates. Data are presented as mean ±SD, as indicated in the figure legends. Statistical tests were selected according to the experimental design. For comparisons between two groups, unpaired two-tailed Student’s *t*-test was used, while comparisons involving multiple groups were performed using one-way ANOVA. A *p* value < 0.05 was considered statistically significant.

### circRNA sequence comparison and candidate filtering

#### circRNA sequences and comparison between circAtlas releases

circRNA sequences were downloaded from circAtlas versions 2.0 and 3.0 (https://ngdc.cncb.ac.cn/circatlas/). Because the reverse complement of a structured RNA can also form secondary structures, and no definitive structural signature defines an IRES (52), we considered both sequence orientations. This allows us to account for annotation differences, and therefore matched sequences by exact or partial reverse complementarity rather than by identifier, yielding more than 180,000 cases. We retained the 27,110 circRNAs with a single match between releases. IRESFinder was then applied independently to both sequences of each circRNA, identifying 16,355 circRNAs with at least one high-scoring IRES-like element in each release.

#### circRNAs regulated by SAM68

The list of circRNAs from circAtlas v2.0 that are up- or downregulated upon *Sam68* knockout at three time points of mESC differentiation (day 0, day 3 and day 10) was retrieved from our previous work (9). Intersection of this list (63 circRNAs) with the 16,355 circRNAs retained above yielded three candidates.

#### Prediction of circRNA–RBP interactions

Interactions were predicted with the *cat*RAPID algorithm, which estimates binding potential from van der Waals, hydrogen-bonding and secondary-structure propensities of the protein and RNA sequences (53). *cat*RAPID was used to estimate the binding propensity of circRNAs for murine SAM68 and for a set of translation-related RBPs in mouse and human. RBPs were retrieved from UniProt (release 2026_03) applying the following criteria: reviewed status; annotation with translation-related functions (GO:0006412); RNA-binding activity (GO:0003723); exclusion of ribosomal proteins (GO:0044391); and exclusion of translational repressors (GO:0030371). This yielded 83 mouse and 108 human RBPs, restricting the analysis to proteins that promote translation without belonging to the ribosomal machinery or acting as repressors.

#### Prediction of IRES elements

IRES elements were predicted using IRESFinder, a tool also employed by circATLAS (52). We used mode 2 with default parameters to fragment the sequences and identify candidate regions. Considering the author’s guidelines, fragments with a score > 0.5 were considered indicative of the presence of an IRES element, and fragments with a score < 0.5 of its absence.

## Data availability

Available upon reasonable request to the corresponding author.

## Acknowledgements

We thank Prof. Stefano Gustincich for providing the pRUF plasmid and for helpful discussions. We thank all members of Tartaglia’s lab for insightful discussions. We thank Dr. Davide Mariani for providing the FUS^P525L^ neuroblastoma cell line and for assistance with the immunofluorescence protocol, and Dr. Lorenzo Stufera Mecarelli for advice on the analysis of the immunofluorescence images. We also acknowledge the members of the RNA Technologies Flagship @IIT for their constructive input.

