## Supplementary Information for "A circRNA-derived IRES-like element enables programmable translational activation of the endogenous chaperone HSP90"

### Supplemental information

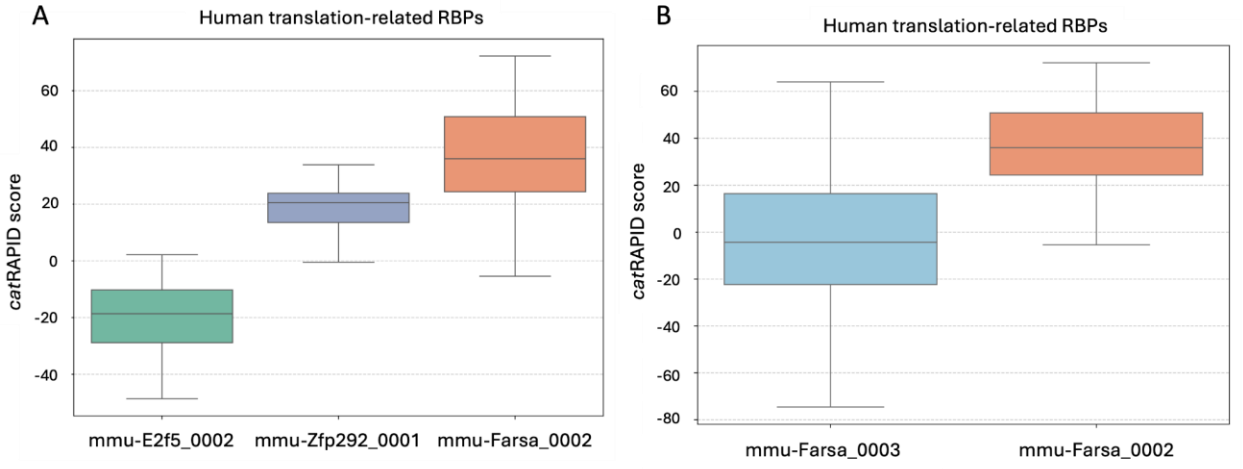


**Figure S1**: **Identification of *mmu-Farsa_0002* as a circRNA-derived IRES-like element, relevant to Figure 1.** A: Propensity to interact with human translation-related RBPs of the three circRNAs candidates (*mmu-Farsa_0002, mmu-Zfp292_0001* and *mmu-E2f5_0002*). On the y axis the interaction propensity score calculated using *cat*RAPID is shown for each RBP. B: *mmu-Farsa_0002* (v2.0) and *mmu-Farsa_0003* (v3.0) propensities to interact with human translation-related RBPs. On the y axis the interaction propensity score calculated using *cat*RAPID is shown for each RBP.

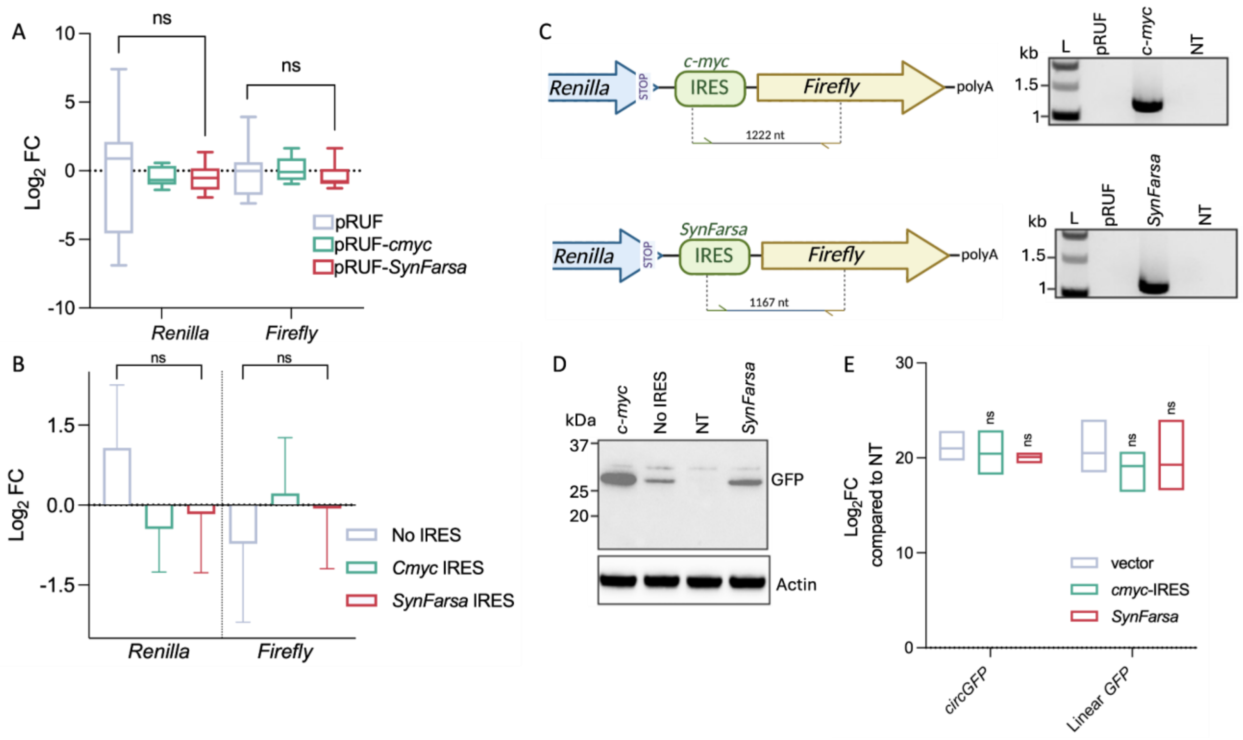


**Figure S2: SynFarsa exhibits an IRES-like activity in cis, relevant to Figure 2.** A and B: real-time PCR analysis demonstrating constant RNA levels of *Renilla* and *Firefly* upon transfection of pRUF, pRUF-*cmyc* and pRUF-*SynFarsa*. Box plots show the median line and data range derived from 8 biological replicates. Statistical non-significance was assessed via unpaired t-test ($p<0.05$). C: schematic illustration of the primer design for RT-PCR, verifying the expression of the IRES of interest following transfection of the *in vitro* transcribed bicistronic RNAs containing *cmyc* IRES or *SynFarsa* IRES (on the left). Agarose gel images depict the PCR products observed in the presence of *cmyc* IRES or *SynFarsa* IRES, while no products are detected in the negative control samples lacking an IRES (pRUF) or those that were not transfected (NT) (on the right). D: Western blot analysis to verify the expression of GFP in the presence of *cmyc* IRES, No IRES, untransfected cells (NT) and *SynFarsa* IRES. Actin was used as a loading control. E: Real-time PCR analysis, demonstrating that transfection of plasmids producing circularizing GFP (*circGFP*) fused with *cmyc* IRES or *SynFarsa* IRES yields consistent expression levels of circular and linear *GFP* expression when compared to the empty plasmid (vector). Box plots depict the median line and data range, sourced from 3 biological replicates. Statistical significance was assessed via two-way ANOVA ($p<0.05$).

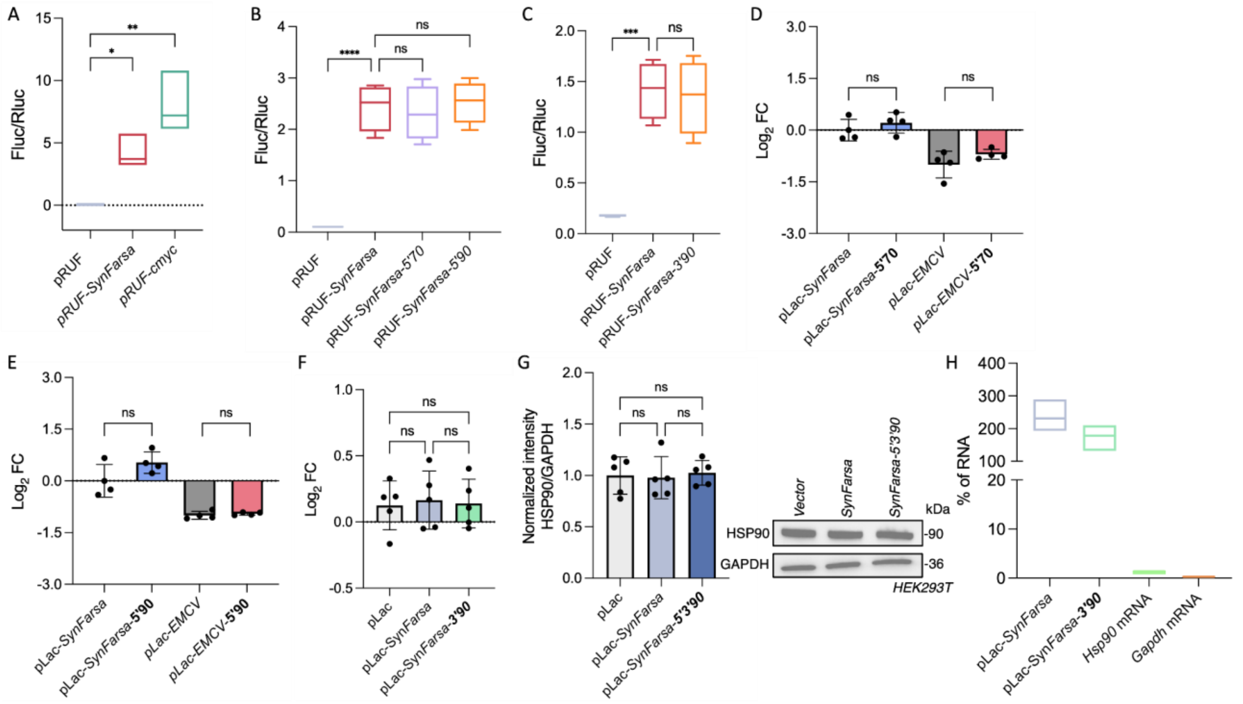


**Figure S3: The *SynFarsa* model for targeted translational activation of endogenous mRNAs, relevant to Figure 3. A:** Dual-luciferase assay performed 24 hours post-transfection of HEK293T cells to confirm *SynFarsa* activity in human cells. *Fluc* activity was normalized to *Rluc* and the impact of each tested sequence on *Fluc* was assessed by comparing normalized *Fluc/Rluc* ratios to that of pRUF control. Box plots represent data from 4 independent biological replicates and statistical significance was assessed by ordinary one-way ANOVA ($p<0.05$). **B** and **C**: Dual-luciferase assay performed 24 hours post-transfection of HEK293T cells to confirm that fusing gRNA sequences to the IRES did not compromise its activity. *Fluc* activity was normalized to *Rluc* and the impact of each tested sequence on *Fluc* was assessed by comparing normalized *Fluc/Rluc* ratios to that of pRUF-*SynFarsa* control. Box plots represent data from 4 independent biological replicates and statistical significance was assessed by ordinary one-way ANOVA ($p<0.05$). **D** and **E**: Real-time PCR analysis showing unchanged RNA levels of *hsp70* (**D**) and *hsp90* (**E**) mRNAs after targeting their 5'UTRs. Box plots represent data from 4 independent biological replicates and non-significance was assessed through t-test ($p<0.05$). **F**: Real-time PCR analysis showing unchanged RNA levels of *hsp90* mRNA after targeting its 3'UTR. Box plots represent data from 5 independent biological replicates and non-significance was assessed through t-test ($p<0.05$). **G**: Western blots showing no change in HSP90 protein levels in HEK293T cells following 48 hours of transfection of *SynFarsa-5'3'90* construct. Quantification plots normalized to the loading control GAPDH are shown on the left, and representative plots are shown on the right. N= 5 biological replicates and non-significance was assessed through one-way ANOVA ($p<0.05$). **H**: Validation of the circular conformation of the overexpression vector pLac by RNase R treatment and RT-qPCR analysis. Circular RNAs are resistant to degradation by RNase R but linear transcripts were efficiently degraded. Levels of RNA are shown as percentage of remaining RNA compared to untreated controls, for 3 independent biological replicates.

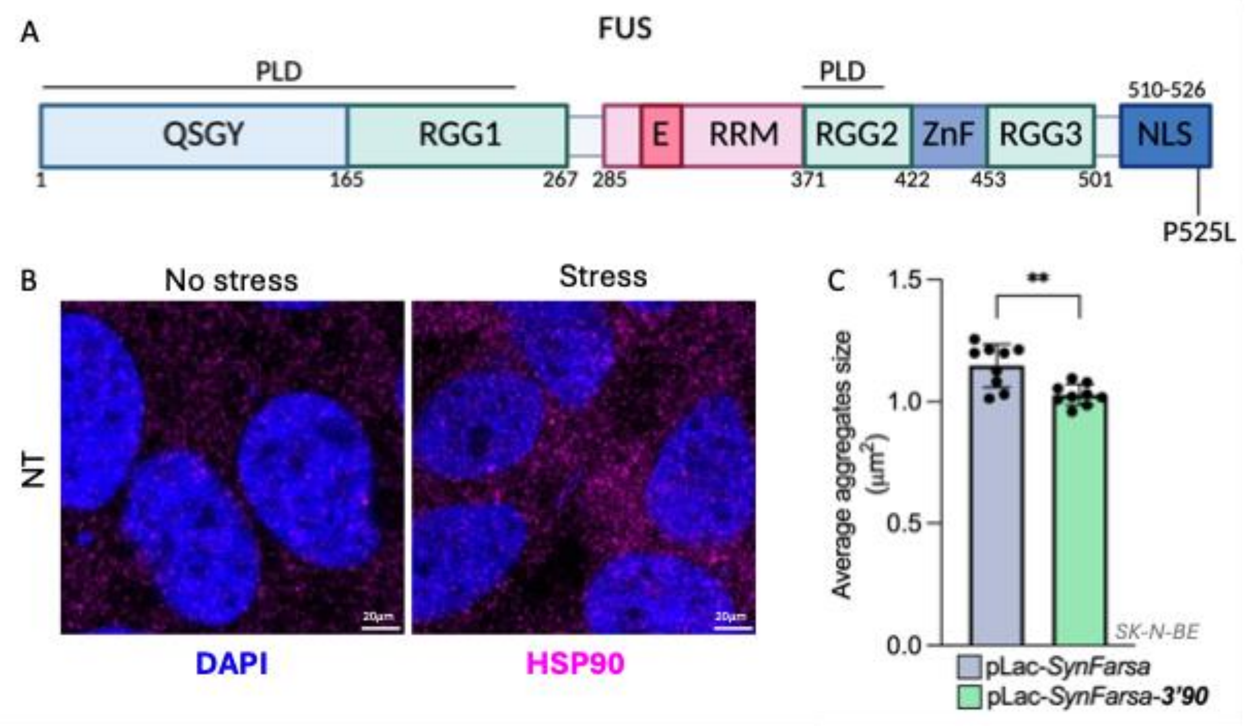


**Figure S4: *SynFarsa*-mediated HSP90 upregulation modulates FUS aggregation, relevant to Figure 4. A**: Schematic representation of human FUS protein and its domains, showing the localization of the P525L mutation. QSGY and RGG, regions rich in the stated amino acids; PLD, prion-like domain; E, nuclear export signal; RRM, RNA recognition motif; ZnF, zinc finger domain; NLS, nuclear localization sequence. Numbers indicate amino acid positions. **B**: representative confocal images of $FUS^{P525L}$-expressing SK-N-BE cells showing HSP90 fluorescence intensity in stress condition compared to non-stressed cells. DAPI staining and HSP90 fluorescence signal are shown. Scale bar = 20 µm. **C**: average aggregates size quantification (preliminary data). Bar plots represent values from 9 technical replicates. Significance was assessed using unpaired t-test ($p<0.05$).

### Supplementary tables

| Plasmid | Source |
|---|---|
| pRUF | Prof. Gustincich lab (87) |
| pCDNA3.1(+)LACCASE2 MCS Exon Vector | Addgene Cat#69893 |
| pcDNA3.1(+) ZKSCAN1 MCS-WT Split GFP | Addgene Cat#69909 |
| pUC57 | Genescript Cat#SD1176 |

*Table 1: Plasmids used in the project*

| Primer | Sequence 5'-3' | Usage |
|---|---|---|
| Cloning in pRUF for luminescence assay | | |
| *SynFarsa*_pRUF_F | CCGCTCGAGCTCGCAGGAAGAAAGTATCATGTTGG | |

| | | |
|---|---|---|
| *SynFarsa*_pRUF_R | GTAGTTCATATGGATCCTGAAAACCTACTGGGTGAGCAAG | Cloning Farsa in pRUF |
| *SynFarsa*_*Frg1*_F | CCGCTCGAGCGGACCTTGAGCAAGGGGTG | Cloning *Frg1* |
| *SynFarsa*_*Frg1*_R | GTAGTTCATATGGATCCTGAAAACCTACTGGGTGAGCAAG | |
| *SynFarsa*_*Frg2*_F | CCGCTCGAGGGGGTGCAGGTGGCCACTAT | Cloning *Frg2* |
| *SynFarsa*_*Frg2*_R | GTAGTTCATATGACTGGGTGAGCAAGGGCAAGG | |
| Neg_Ctrl_F | CCGCTCGAGTGAGTTTTTGTTCTTAGATAACCCACATTAGATGA | Cloning Neg control |
| Neg_Ctrl_R | GTAGTTCATATGGACAACTCTTATAAAAAGCATCCAAGCACA | |
| SV40_del_F | AAGCTTGATTCTTCTGACACAACAG | SV40 deletion |
| SV40_del_R | AAGGTACCTAACCAAGTTCCTCTT | |
| *SynFarsa*_5'70_F | AGGATCGCCTTCCTGGCTCCATATGCCATGGAAGACGCCAAAAA | Cloning 5'70 |
| *SynFarsa*_5'70_R | GGCATATGGAGCCAGGAAGGCGATCCTGAAAACCTACTGG | |
| *SynFarsa*_5'90_F | GGATCTCGCAGGGCCACACATATGCCATGGAAGACGCCAAAAAC | Cloning 5'90 |
| *SynFarsa*_5'90_R | GGCATATGTGTGGCCCTGCGAGATCCTGAAAACCTACTGG | |
| *SynFarsa*_3'90_F | GGATCATAGCATTTCGACAAGTTTTTATTGAGTCTCATATGCCATGGAAG | Cloning 3'90 |
| *SynFarsa*_3'90_R | ATATGAGACTCAATAAAAACTTGTCGAAATGCTATGATCCTGAAAACCTA | |
| Cloning in pcDNA3.1(+) ZKSCAN1 MCS-WT Split GFP for split GFP assay | | |
| Lin_pZKSCAN_F | TATGGCGTTAACCACCATGGC | Linearization of pZKSCAN |
| Lin_pZKSCAN_R | CGTTAACGACTCGGAACGGTTAC | |
| *SynFarsa*_pZKSCAN_F | ACCGTTCCGAGTCGTTAACGCTCGCAGGAAGAAAGTATC | Cloning *SynFarsa* |
| *SynFarsa*_pZKSCAN_R | CCATGGTGGTTAACGCCATAGATCCTGAAAACCTACTG | |
| *Cmyc*_pZKSCAN_F | ACCGTTCCGAGTCGTTAACGTAATTCCAGCGAGAG | Cloning c-myc |
| *Cmyc*_pZKSCAN_R | CCATGGTGGTTAACGCCATACGTCGCGGGAGGCTG | |
| Empty_pZKSCAN_F | TCCGAGTCGTTAACGTATGGCGTTAACCACCATGG | Empty pZKSCAN |
| Empty_pZKSCAN_R | GTGGTTAACGCCATACGTTAACGACTCGGAACGGTTA | |
| Cloning in pCDNA3.1(+) LACCASE2 MCS Exon Vector for overexpression | | |
| Lin_pLac_R | TCTGCATAAAATAAAAAAAATATATATAACAATAAAAC | Linearization of pLac |
| Lin_pLac_F | GTAAGTATTCAAAATTCCAAAATTTTTT | |
| *SynFarsa*_pLac_F | TTTTTTTTATTTTATGCAGACTCGCAGGAAGAAAGTATCATGTTGGT | Cloning *SynFarsa* |
| *SynFarsa*_pLac_R | TTGGAATTTTGAATACTTACGATCCTGAAAACCTACTGGGTGAGC | |
| pLac_5'70_F | CAGGATCGCCTTCCTGGCTCGTAAGTATTCAAAATTCCAAAATT | |

| | | |
|---|---|---|
| pLac_5’70_R | TACTTACGAGCCAGGAAGGCGATCCTGAAAACCTACTGGGTGAG | Cloning *SynFarsa*_5’ 70 |
| pLac_5’90_F | CAGGATCTCGCAGGGCCACAGTAAGTATTCAAAATTCCAAAATT | Cloning *SynFarsa*_5’ 90 |
| pLac_5’90_R | TACTTACTGTGGCCCTGCGAGATCCTGAAAACCTACTGGGTGAG | |
| pLac_EMCV_F | TTTTTTTTATTTTATGCAGACCCCTCTCCCTCCCCCCCC | Cloning EMCV in pLac |
| pLac_EMCV_R | AAAAAATTTTGGAATTTTGAATACTTACTTATCATCGTGTTTTTCAA AGG | |
| EMCV_5’70_F | ATAAGCCTTCCTGGCTCGTAAGTATTCAAAATTCCAAAAT | Cloning EMCV_5’70 |
| EMCV_5’70_R | ACTTACGAGCCAGGAAGGCTTATCATCGTGTTTTTCAAAG | |
| EMCV_5’90_F | ATAATCGCAGGGCCACAGTAAGTATTCAAAATTCCAAAAT | Cloning EMCV_5’90 |
| EMCV_5’90_R | ACTTACTGTGGCCCTGCGATTATCATCGTGTTTTTCAAAG | |
| *SynFarsa*_3’70_F | ATATATTTTTTTTATTTTATGCAGACTCGCAGGAAGAA | Cloning *SynFarsa*_3’70 from pUC57 |
| *SynFarsa*_3’70_R | AAATTTTGGAATTTTGAATACTTACTGAATTTTTTACCGG | |
| *SynFarsa*_3’90_F | ATATATTTTTTTTATTTTATGCAGACTCGCAGGAAGAA | Cloning *SynFarsa*_3’90 from pUC57 |
| *SynFarsa*_3’90_R | AAATTTTGGAATTTTGAATACTTACAGACTCAATAAAAAC | |
| *SynFarsa*_5’3’70_F | ATATATTTTTTTTATTTTATGCAGACTCGCAGGAAGAA | Cloning *SynFarsa*_53’ 70 from pUC57 |
| *SynFarsa*_5’3’70_R | AAATTTTGGAATTTTGAATACTTACTGAATTTTTTACCGG | |
| *SynFarsa*_5’3’90_F | ATATATTTTTTTTATTTTATGCAGACTCGCAGGAAGAA | Cloning *SynFarsa*_53’ 90 from pUC57 |
| *SynFarsa*_5’3’90_R | AAATTTTGGAATTTTGAATACTTACAGACTCAATAAAAAC | |

*Table 2: Primers used in the project. Highlighted in bleu (forward) and green (reverse) the complementary region with the targeted mRNA*

| | **Gene reference** |
|---|---|
| *Farsa* | mmu-Farsa_0002, circAtlas (Version 2.0) |
| *Hsp70* | ENST00000375671.7 HSPA1A-201 |
| *Hsp90* | ENST00000216281.13 HSP90AA1-201 |

*Table 3: Gene references of the transcripts used in the project.*

| **Fragment** | **Selected sequence 5’-3’** | **Reverse complement 5’-3’** |
|---|---|---|

| | | |
|---|---|---|
| 5'UTR of *hsp70* | CGGAAGGACCGAG | GCCTTCCTGGCTC |
| 5'UTR of *hsp90* | AGCGTCCCGGTGT | TCGCAGGGCCACA |
| 3'UTR of *hsp70* | AATACACTTAACTCAGGCCATTTTTTAAGT | TTATGTGAATTGAGTCCGGTAAAAAATTCA |
| 3'UTR of *hsp90* | TATCGTAAAGCTGTTCAAAAATAACTCAGA | ATAGCATTTCGACAAGTTTTTATTGAGTCT |
| Linker | CCACCGCGGTGGCCGCTCTAGATTCATCACTTATTCAGGC | N/A |

*Table 4: Sequences of the UTRs fragments and the linker used in the project.*

| Antibody | Specie | Reference | Application |
|---|---|---|---|
| HSP70 | Rabbit | Cell Signaling, BK4872S | Western blot |
| HSP90 | Rabbit | Cell Signaling, BK4877S | |
| GAPDH | Rabbit | Cell Signaling, BK2118S | |
| HS90 | Rabbit | Cell Signaling, BK4877S | Immunofluorescence |
| Anti-FLAG | Mouse | Merck, F3165-1MG | |
| Anti-rabbit IgG Alexa Fluor 647 | - | Cell Signaling, 4414 | |
| Anti-mouse IgG Alexa Fluor 488 | - | Thermo Fisher Scientific, 488A11029 | |

*Table 5: Antibodies used in the project.*

| Primer | F/R | Sequence |
|---|---|---|
| *Firefly (Fluc)* | F | TCTGAGGAGCCTTCAGGATT |
| | R | AGATGGAACCTCTTGGCAAC |
| *Renilla (Rluc)* | F | TTGAATCATGGGATGAATGG |
| | R | GGACGACGAACTTCACCTTT |
| *Gapdh* (mouse) | F | GCAGTGGCAAAGTGGAGATT |
| | R | GCAGAAGGGGCGGAGATGA |
| *Cmyc* | F | ACGAAACTTTGCCCATAGCAG |
| | R | CAAATGGGCAGAATAGCCT |
| *SynFarsa* | F | CACTATCTGGGAGCACACC |
| | R | GATGATCTCCAGTGGCTCTTG |

| | | |
|---|---|---|
| *Cmyc* expression (bicistronic vector) | F | ACGAAACTTTGCCCATAGCAG |
| | R | AGATGGAACCTCTTGGCAAC |
| *SynFarsa* expression (bicistronic vector) | F | CACTATCTGGGAGCACACC |
| | R | AGATGGAACCTCTTGGCAAC |
| circGFP | F | ATGACCGACAAGGCCAAGAA |
| | R | CAGGTAGTGGTTATCGGGCA |
| linGFP | F | GTAGTGGTTATCGGGCAGCA |
| | R | CATCATGACCGACAAGGCCAA |
| *Hsp70* | F | CCTGTGTTTGCAATGTTGAAATTTTTTGGT |
| | R | ACTTTGAAGTAACCAACTTAAAAAATGGCC |
| *Hsp90* | F | GGAGTATCTGGATGACAGTTAGTGCC |
| | R | GCAGGGGATGCATGTTGTCTGC |
| *Gapdh* (human) | F | GTCTCCTCTGACTTCAACAGCG |
| | R | ACCACCCTGTTGCTGTAGCCAA |
| *EMCV* | F | GTCTGTAGCGACCCTTTG |
| | R | TTGAATACGCTTGAGGAGAGC |
| *Divergent SynFarsa* | F | CCTCCTGCTTAGACACGCTT |
| | R | AGCACGTGACCAACATGATACT |
| Circular *SynFarsa_3'90* | F | TATCTGGGAGCACACCACG |
| | R | ACTTGTCGAAATGCTATGATCCTG |
| *eGFP* | F | AGGAGCGCACCATCTTC |
| | R | GATGCCCTTCAGCTCGAT |

*Table 6: Primers used for RT-qPCR.*
